# Dealing with many correlated covariates in capture-recapture models

**DOI:** 10.1101/097006

**Authors:** Olivier Gimenez, Christophe Barbraud

## Abstract

Capture-recapture models for estimating demographic parameters allow covariates to be incorporated to better understand population dynamics. However, high-dimensionality and multicollinearity can hamper estimation and inference. Principal component analysis is incorporated within capture-recapture models and used to reduce the number of predictors into uncorrelated synthetic new variables. Principal components are selected by sequentially assessing their statistical significance. We provide an example on seabird survival to illustrate our approach. Our method requires standard statistical tools, which permits an efficient and easy implementation using standard software.

## Introduction

Capture-recapture (CR) methods (e.g. Lebreton *et al.* 1992) are widely used for assessing the effect of explanatory variables on demographic parameters such as survival (Pollock 2002). Generally however, complex situations arise where multiple covariates are required to capture patterns in survival. In such situations, one usually favors a multiple regression-like CR modeling framework that is however hampered by two issues: first, because it increases the number of parameters to be estimated, incorporating many covariates results in a loss of statistical power and a decrease in the precision of parameter estimates; second, correlation among the set of predictors – aka multicollinearity – may alter interpretation (see below).

To overcome these two issues, Grosbois *et al.* (2008) recommended to perform a principal component analysis (PCA) on the set of explanatory variables before fitting CR models. PCA is a multivariate technique that explains the variability of a set of variables in terms of a *reduced* set of *uncorrelated* linear combinations of such variables – aka principal components (PCs) – while maximizing the variance (Jolliffe 2002). Grosbois *et al.* (2008) then expressed survival as a function of the PCs that explained most of the variance in the set of original covariates, typically the first one or the first two ones.

However, the main drawback of this approach is that the PCs are selected based on covariates variation pattern alone, regardless of the response variable, and without guarantee that survival is most related to these PCs (Graham 2003). To deal with this issue in the context of logistic regression, Aguilera *et al.* (2006) proposed to test the significance of *all* PCs to decide which ones should be retained, instead of a priori relying on the PCs that explain most of the variation in the covariates.

In this paper, we implement the algorithm proposed by Aguilera *et al.* (2006) to deal with many possibly correlated covariates in CR models, a method we refer to as principal component capture-recapture (P2CR). We apply this new approach to a case study on survival of Snow petrels (*Pagodroma nivea*) that is possibly affected by climatic conditions. In this example, the issue of multicollinearity occurs, and summarizing the set of covariates in a subset of lower dimension is also crucial to get precise survival estimates. Overall, P2CR models can be fitted with statistical programs that perform PCA and CR data analysis. The data and R code are available from GitHub at https://github.com/oliviergimenez/p2cr.

## Methods

We used capture-recapture (CR) models to study open populations over K capture occasions to estimate the probability *ϕ_i_* (*i* = 1, …, K –1) that an individual survives to occasion *i* + 1 given that it is alive at time *i*, along with the probability *p_j_* (*j* = 2, …, K) that an individual is recaptured at time *j* – aka as the Cormack-Jolly-Seber (CJS) model (Lebreton *et al.* 1992). Covariates were incorporated in survival probabilities using a linear-logistic function:

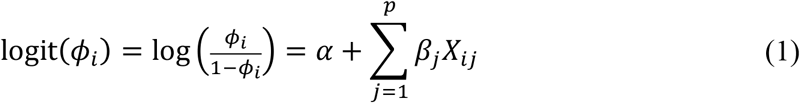

where *α* is the intercept parameter, *X_ij_* is the value of covariate *j* (*j* = 1,…, *p*) in year *i* (*i* = 1,…, K – 1), and *β_j_* is its associated slope parameter. Covariates were standardized to avoid numerical instabilities. To assess the significance of a covariate in CR models, we used the analysis of deviance (ANODEV; Skalski, Hoff & Smith 1993) that compares the amount of deviance explained by this covariate with the amount of deviance not explained by this covariate, the CR model with fully time-dependent survival serving as a reference. The ANODEV test statistic is given by:

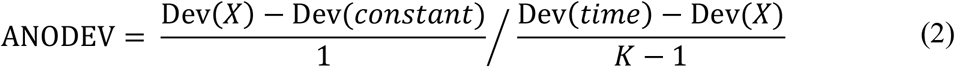

where Dev(*constant*), Dev(*X*) and Dev(*time*) stand for the deviance of models with constant, covariate-dependent and time-dependent survival probabilities. To obtain the associated p-value, the value of the ANODEV is compared with the quantile of Fisher-Snedecor distribution with 1 and K-1 degrees of freedom.

To reduce the dimension of the set of covariates (X_1_, …, X_p_), we used PCA which aims at finding a small number of linear combinations of the original variables – the principal components (PCs) – while maximizing the variance in (X_1_, …, X_p_). Because the variables measurement units often differ, we performed the PCA on the correlation matrix (Jolliffe 2002). To select PCs, we used a forward model selection algorithm as proposed by Aguilera *et al.* (2006) for the logistic regression. The forward algorithm begins with no covariates in the model. Each PC is incorporated in simple linear regression-like CR models and the ANODEV p-value calculated. The PC that has the lowest p-value is added to the null model, say PC_k_. Then the PCs that were not retained are incorporated along with PC_k_ in multiple regression-like CR models, and ANODEV p-values are calculated. In other words, we need to assess the effect of PC_j_ for j ≠ k in the presence of PC_k_ to decide whether PC_j_ should be retained. To do so, Dev(*constant*) and Dev(*X*) are replaced by Dev(PC_k_) and Dev(PC_k_ + PC_j_) in Equation 2, where Dev(PC_k_ + PC_j_) is the deviance of the model with survival as a function of both principal components PC_k_ and PC_j_. We repeat the process until no remaining PC is selected.

All models were fitted using the maximum-likelihood method using MARK (White & Burnham 1999) called with R (Laake 2013).

## Case study

The Snow petrel is a medium sized Procellariiform species endemic to Antarctica that breeds in summer. Birds start to occupy breeding sites in early November, laying occurs in early December and chicks fledge in early March. This highly specialized species only forages within the pack-ice on crustaceans and fishes. Data on survival were obtained from a long-term CR study on Ile des Pétrels, Pointe Géologie Archipelago, Terre Adélie, Antarctica. We refer to Barbraud *et al.* (2000) for more details about data collection. We removed the first capture to limit heterogeneity among individuals, and worked with a total of 604 female capture histories from 1973 to 2002.

The following covariates were included to assess the effect of climatic conditions upon survival variation: sea ice extent (SIE; http://nsidc.org/data/seaice_index/); air temperature, which was obtained from the Météo France weather station at Dumont d'Urville, as a proxy for sea surface temperature; southern Oscillation Index (SOI) as a proxy for the overall climate condition (https://crudata.uea.ac.uk/cru/data/soi/). These environmental variables were averaged over seasonal time periods corresponding to the chick rearing period (January to March: summer period), the non-breeding period (April to June: autumn and July to September: winter), and the laying and incubation period of the same year (October to December: spring). In total, 9 covariates were included in the analysis: sea ice extent in summer (SIEsummer), in autumn (SIEautumn), in winter (SIEwinter), in spring (SIEspring), annual SOI, air temperature in summer (Tsummer), in autumn (Tautumn), in winter (Twinter) and in spring (Tspring).

## Results

The CJS model poorly fitted the data (χ^2^ = 221.2, df = 127, p << 0.01), and a closer inspection of the results revealed that the lack of fit was explained by a trap-dependence effect (Test2CT, χ^2^ = 103.1, df = 27, p << 0.01). Consequently, we estimated two recapture probabilities that differed according to whether or not a recapture occurred the occasion before (Pradel 1993). By first attempting to simplify the structure of recapture probabilities, we were led to consider an additive effect of time and a trap effect (Supplementary material). Estimates of recapture probabilities ranged from 0.14 (standard error [SE] = 0.07) to 0.79 (SE = 0.09) when no recapture occurred the occasion before and from 0.25 (SE = 0.18) to 0.89 (SE = 0.09) when a recapture occurred the occasion before (Supplementary material).

Because of multicollinearity, we were led to counterintuitive estimates of regression parameters in the CR model including all covariates (Supplementary material): the coefficient of SIE in autumn was estimated at 0.5 (SE = 0.24) and that of SIE in winter was estimated at –0.5 (SE = 0.21) while these two covariates were significantly positively correlated (r = 0.67, p < 0.01).

When we applied the P2CR approach, the algorithm selected two PCs, namely PC3 (F_1,27_ = 7.34, p = 0.01) at step 1 and PC4 (F_1,26_ = 4.63, p = 0.04) at step 2 (Supplementary material), but never did we pick PC1 as we would have done using a classical approach (Grosbois et al. 2008). PC3 was positively correlated to SIE in summer and negatively correlated to temperature in winter, while PC4 was positively correlated to temperature in spring and negatively correlated to SIE in summer (Supplementary material). Survival increased with increasing values of PC3 (Figure 1), with high values of SIE in summer and low values of temperature in winter (resp. low values of SIE in summer and high values of temperature in winter) corresponding to high (resp. low) survival.

**Figure 1.**
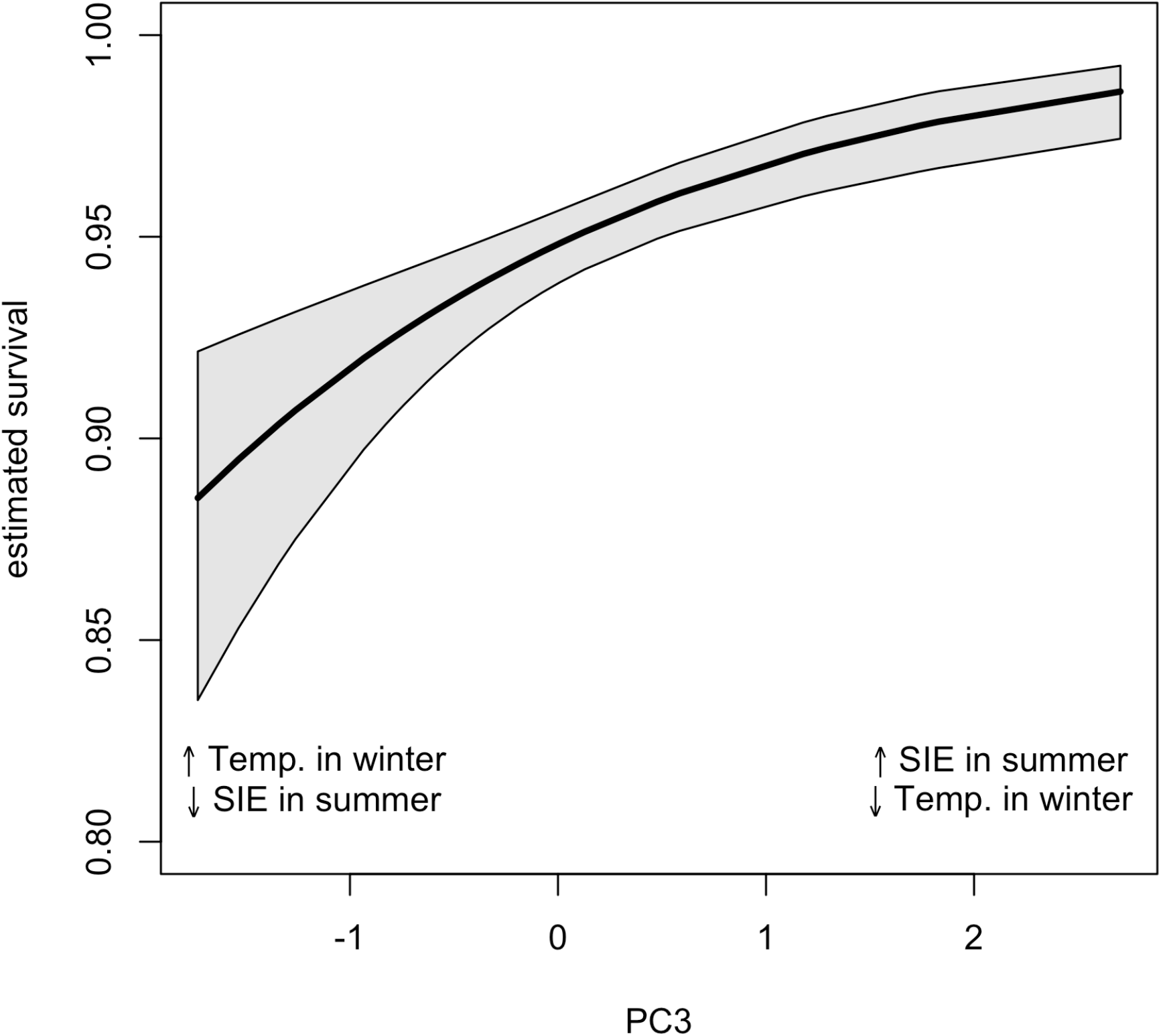
Survival of Snow petrel as a function of PC3.

Survival decreased with increasing values of PC4 (Figure 2), with high values of temperature in spring and low values of SIE in summer (resp. low values of temperature in spring and high values of SIE in summer) corresponding to low (resp. high) survival.

**Figure 2.**
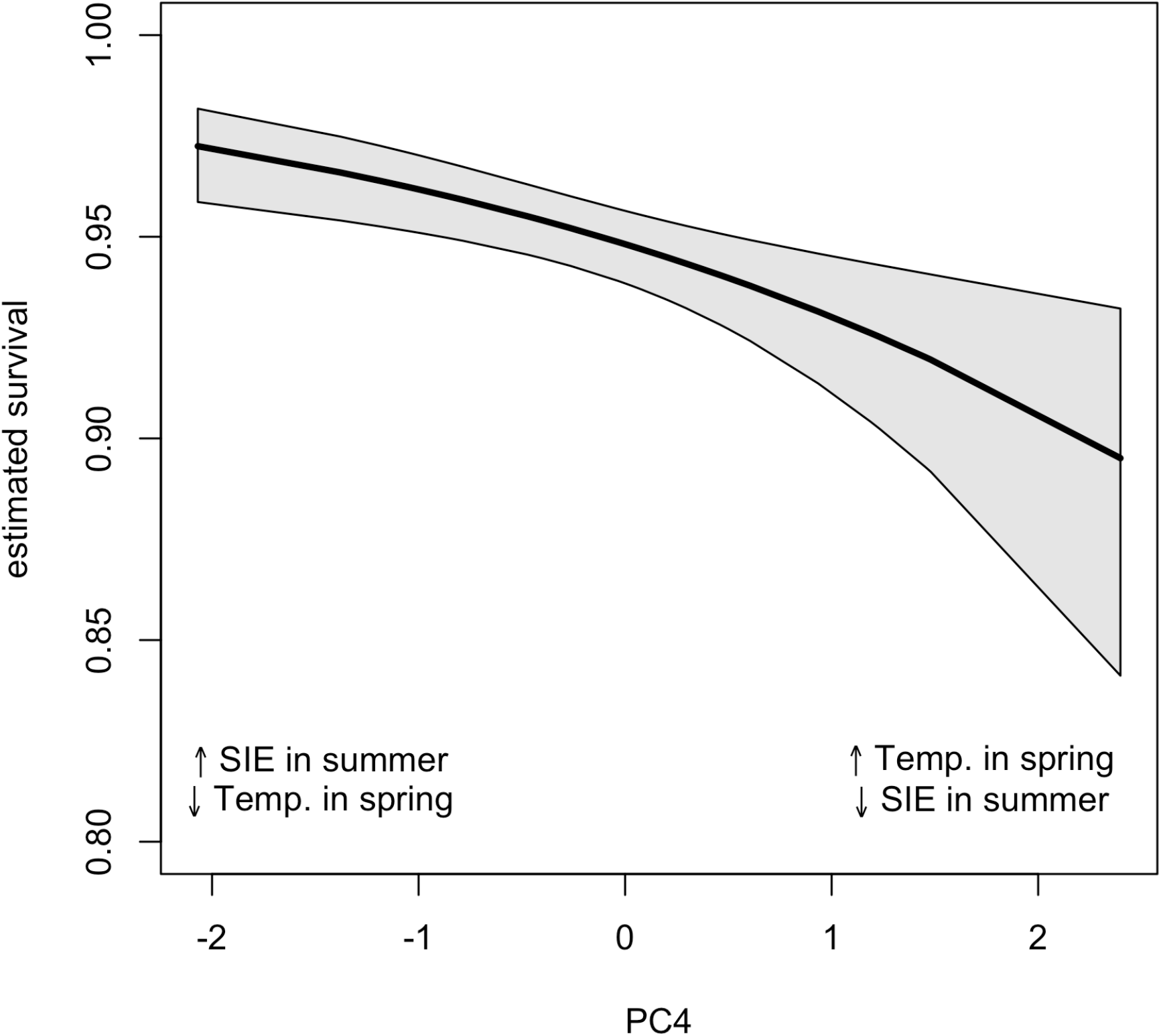
Survival of Snow petrel as a function of PC4.

The P2CR approach also led to more precise survival estimates when compared to the model incorporating all original covariates (Figure 3).

**Figure 3.**
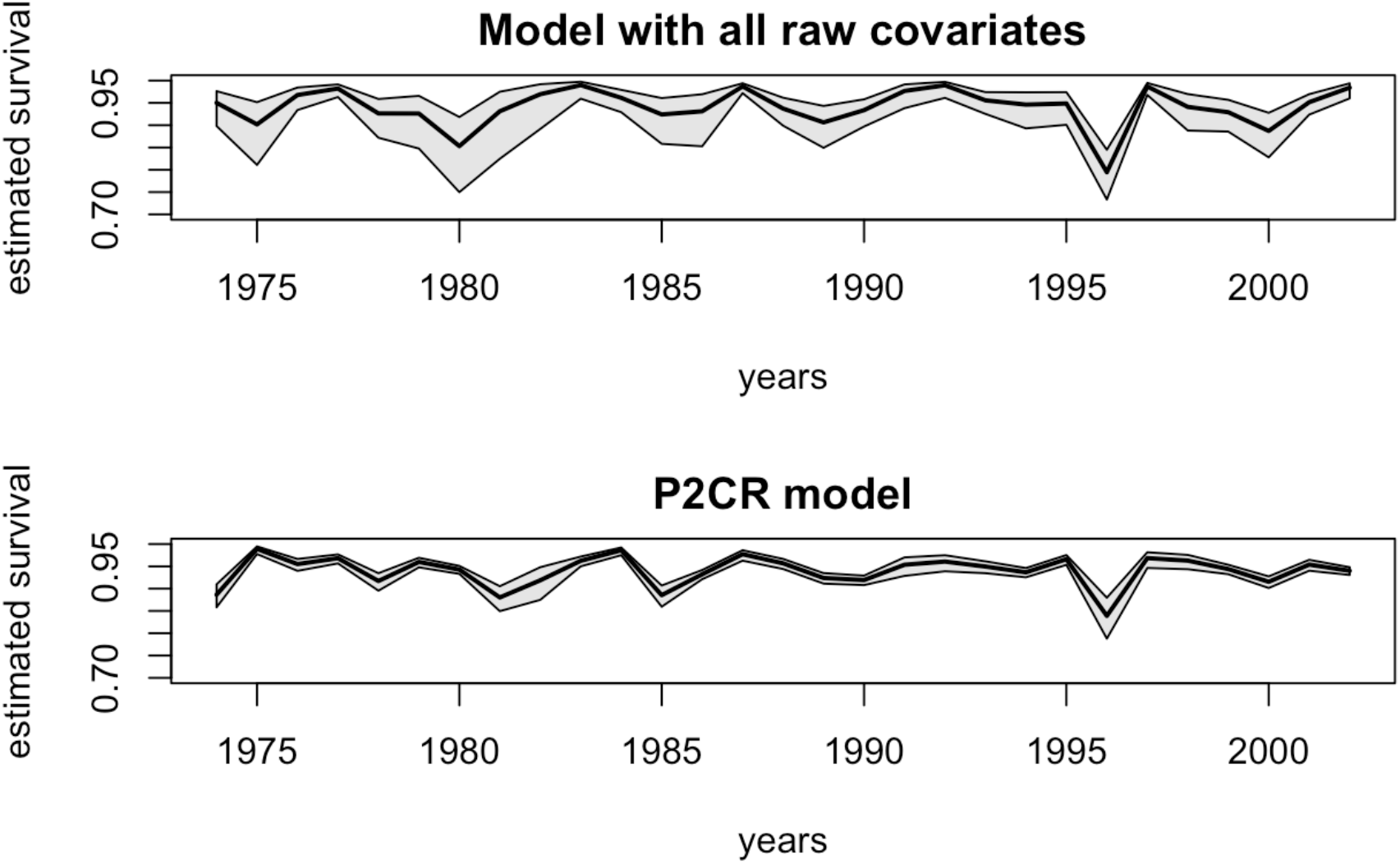
Survival of Snow petrel over time as estimated from the model with all original covariates (top panel) vs. the PC2R model (bottom panel).

## Discussion

We introduce a new approach combining principal component analysis and capture-recapture models to deal with many possibly correlated explanatory covariates. Our approach requires standard statistical tools, which allows an efficient and easy implementation using standard software.

### Snow petrels and climatic conditions

In summer, snow petrels exclusively forage within the pack-ice tens to hundreds of kilometers from the colony where they catch sea ice-associated species, such as Antarctic silverfish (*Pleuragramma antarcticum*) and Euphausiids, to feed their chick (Ridoux & Offredo 1989). This is an energetically demanding period for breeding adults and, during years with reduced sea-ice extent, food resources may be less abundant and snow petrels may be forced to cover larger distances to find suitable foraging habitats, with potential survival costs. Assuming air temperature was a proxy of sea surface temperature variations, the negative effect of warmer temperatures on survival is coherent with general patterns found between sea surface temperature and demographic parameters in seabirds (Barbraud *et al.* 2012). In many marine ecosystems warmer temperatures are associated with decreased primary production and food resources for top predators. Although the low survival in 1996 corresponded to a year with reduced sea-ice extent in summer, the drop in survival was high and remains unexplained at the moment.

### Principal component CR models

When multiple covariates have to be considered to estimate survival, both issues of dimensionality and multicollinearity can lead to biased estimates, inflated precision as well as lack of statistical power. In such a context, the P2CR modeling framework has proved particularly useful in our example, mainly because few PCs were selected which were easily interpretable. We acknowledge that PCs with little interpretability might have been picked up by our method. To make the interpretation easier, PCA results can be post-processed by rotating axes to improve correlations between raw variables and PCs like in the varimax method (Kaiser 1958). Recent developments in the field of multivariate analyses could also be useful, like methods to handle with missing values in PCA (Dray & Josse 2015).

In statistical ecology, one of our objectives is to try and explain variation in state variables such as abundance, survival and the distribution of species. Dimension-reduction methods are promising to deal with many correlated covariates for the analysis of CR or occupancy data.

## Acknowledgements

We greatly acknowledge all of the wintering fieldworkers involved in the monitoring programs in Terre Adélie since 1962, and Dominique Besson for the management of the database. The study on petrels was supported by Expéditions Polaires Françaises, Institut Paul Emile Victor (program IPEV 109, resp. H. Weimerskirch) and Terres Australes et Antarctiques Françaises.

